# The Unfolded Protein Response Sensor PERK Mediates Mechanical Stress-induced Maturation of Focal Adhesion Complexes in Glioblastoma Cells

**DOI:** 10.1101/2024.03.04.583281

**Authors:** Mohammad Khoonkari, Dong Liang, Marleen Kamperman, Patrick van Rijn, Frank A.E. Kruyt

**Affiliations:** Department of Medical Oncology, University of Groningen, University Medical Center Groningen, Hanzeplein 1, 9713 GZ, Groningen, the Netherlands; Zernike Institute for Advanced Materials, University of Groningen, Nijenborgh 4, 9747 AG Groningen, the Netherlands; Department of Biomedical Engineering-FB40, University of Groningen, University Medical Center Groningen, A. Deusinglaan 1, 9713 AV Groningen, the Netherlands; W.J. Kolff Institute for Biomedical Engineering and Materials Science-FB41, Groningen, University of Groningen, University Medical Center Groningen, A. Deusinglaan 1, 9713 AV Groningen, the Netherlands

**Keywords:** Glioblastoma, extracellular matrix stiffening, mechanical stress, PERK, focal adhesion complex

## Abstract

Stiffening of the brain extracellular matrix (ECM) in glioblastoma leads to mechanical stress, which is known to contribute to tumor formation and progression. Previously, we found that protein kinase R (PKR)-like endoplasmic reticulum kinase (PERK), a component of the unfolded protein response (UPR), plays a role in the adaptation of glioblastoma stem cells (GSCs) to matrix stiffness through PERK/FLNA dependent F-Actin remodeling. Here, we found that increases in matrix stiffness induces differentiation of GSCs that was not seen in PERK deficient GSCs. Furthermore, we investigated whether PERK is involved in detecting changes in matrix stiffness through focal adhesion complex (FAC) formation and maturation, which are instrumental for transmitting ECM dependent signaling. In PERK deficient GSCs, Vinculin and Tensin expression was decreased, while Talin and Integrin-β1 expression was kept at the same level compared to PERK proficient cells. In addition, in the absence of PERK, Tubulin expression is sharply increased coupled with low Vimentin expression, which was observed as an opposite trend in the presence of PERK. In conclusion, our study reveals a novel role for PERK in regulating the formation of FACs during matrix stiffening, possibly associated with its regulatory capacity in F-Actin remodeling.

## Introduction

Glioblastoma is the most aggressive adult brain tumor with a high mortality rate [1]. Despite multimodal treatment with surgery, chemotherapy, and radiotherapy, the median overall survival rates remain less than 2 years. Glioblastomas originate most often within the cerebral cortex and have a high propensity to invade other parts of the brain. This invasive nature hampers surgical resection, and together with inherent therapy resistance of glioblastoma cells, leads to frequent tumor recurrence and rapid disease progression [2,3]. Tumor heterogeneity is considered to be a main cause of therapy resistance in glioblastoma, in which cancer stem cells (CSCs) are thought to play an important role [4,5]. Glioblastoma stem cells (GSCs) have been identified as highly malignant cells that drive tumor growth and progression [6]. These cells possess self-renewal capabilities, exhibit a strong capacity for tumor initiation, and demonstrate high plasticity, which contributes to their aggressive nature. GSCs and cellular plasticity are strongly regulated by the tumor microenvironment (TME) [7].

Recently, it has been recognized that in addition to biological cues, physical abnormalities also contribute to aggressive tumor behavior such as rapid proliferation, metastasis, and therapy resistance [8,9]. Among the physical traits of cancer [10], stiffness is known as a major accelerator of glioblastoma tumor formation and progression [11,12]. Normal soft brain extracellular matrix (ECM) with stiffness of around 1 KPa, undergoes stiffening, reaching approximately 40 kPa due to the overexpression of ECM components [13], particularly hyaluronic acid (HA) and proteoglycans [11,14–16]. These alterations lead to changes in the physicochemical and mechanical properties of the brain tissue, triggering multiscale cellular adaptations in glioblastoma cells through mechanotransduction signaling pathways [10,13,17,18].

The focal adhesion complex (FAC) serves as a central hub for cellular mechanosensing. These dynamic protein complexes enable the connection between the cytoskeleton of cells and the ECM [19–21]. Focal adhesions (FAs) are in a state of constant flux, with proteins continuously associating and dissociating to transmit signals throughout the cell, impacting various processes ranging from cell motility to the cell cycle [22,23]. FAs act as sensors capable of detecting changes in the structure and properties of the ECM, thereby initiating adaptive cellular responses. The interaction between FAs and ECM primarily involves Integrins [24–26]. Integrins bind to extracellular proteins via short amino acid sequences, such as the RGD motif [27]. FAs can disassemble or mature into larger and stable FACs by recruiting additional proteins such as Talin, Paxillin, Vinculin, and Tensin that further promote integrin clustering and establish links between the FAC and the actin cytoskeleton [18,28–30]. The recruitment of these components to FAs occurs in an ordered and sequential manner, leading to the formation of mature and stationary ECM interacting FACs that actively transmit signals [21,26,31].

While various proteins contribute to the construction of focal adhesion complexes (FACs), not all of them are directly involved in mechanotransduction. There is a specific subcomplex within FACs that plays a direct role in cell-ECM binding dynamics, force transmission, and the active signaling of mechanical stress, leading to cellular adaptations. This subcomplex includes Tensin, Vinculin, Talin, and Integrin-β1. These proteins work together to mediate the dynamic interactions between the cell and the ECM, transmit forces, and initiate signaling pathways in response to mechanical stress [20,23,32,33]. The first step of FACs formation is the binding of Talin with Integrins, known as the backbone of FACs [30]. Subsequently, Vinculin and Tensin are recruited to reinforce FAC protein assembly [22,34]. Notably, Vinculin plays a significant role in the maturation of FACs by binding to actin filaments. Once the connection between FACs and the actin cytoskeleton is established, signal transduction is activated, initiating cytoskeleton remodeling as part of the cellular adaptive response to mechanical stress. Although cytoskeleton remodeling and cellular adaptation to matrix stiffness is mostly regulated by the F-Actin network, microtubules (Tubulin), and intermediate filaments such as Vimentin also play a role. Both Vimentin and Tubulin interact with FACs and can mediate mechano-adaptive responses that involve interactions with F-Actin and Filamin-A (FLNA).

Recently, we reported that the endoplasmic reticulum (ER) stress and unfolded protein response (UPR) sensor PKR-like ER kinase (PERK) mediates an adaptive cellular response in GSCs towards increasing substrate/ECM stiffness mimicked by stiffness tunable hydrogels [35]. PERK is known to have both a kinase function and a protein scaffold function by which it regulates a number of cellular processes that have been related either to restoring proteostasis, or to regulate F-Actin remodeling in order to facilitate ER – cell membrane interaction for maintaining calcium homeostasis [36,37]. The latter involves PERK-FLNA interactions, FLNA being a regulator of F-Actin remodeling, a mechanism which we recently identified to be involved in stiffness dependent increases in F-Actin polymerization in GSCs, which was associated with cell elongation and increased cell proliferation and migration [35,36]. However, the potential involvement of PERK in sensing alterations in ECM stiffening by GSCs remains unknown.

Here, we examined if PERK is connected to FAC formation and thus in sensing and mediating cellular responses to alterations in ECM stiffness. Human blood plasma/alginate hydrogels with tunable stiffness were used to mimic brain ECM stiffening in GB [35]. The expression of different components of the FAC in patient-derived GSCs was studied in relation to substrate stiffness. In addition to the actin cytoskeleton, also the involvement of microtubules (Tubulin) and the intermediate filaments (Vimentin) was investigated for sensing capabilities and adapting to mechanical stress. We found PERK to be required for FAC maturation in particular for the recruitment of Vinculin and Tensin. Stiffness dependent increases in Vimentin expression were also affected by PERK. In the absence of PERK, at lower matrix stiffness ranges, Tubulin expression increased, suggesting a compensatory mechanism for impaired F-Actin remodeling and cellular adaptation to mechanical stress.

## Materials and methods

### Preparation of stiffness tunable hydrogel

Human blood plasma (HBP)/Alginate hydrogels were generated as described earlier [35]. In brief, three different concentrations of Alginate (0.2, 0.9 and 1.8 w/v %) were mixed with HBP for generating hydrogels with different stiffnesses of 1, 12 and 35 kPa, respectively. HBP was diluted 1:1 by adding Neurobasal medium (NBM) with 2% B27 supplement, 20 ng/ml bFGF, 20 ng/ml EGF and 1% L-glutamine and FBS 10% (All purchased from Sigma-Aldrich, Darmstadt, Germany). The hydrogels stiffness was checked using a rheometer as described before [35].

### Cell culture

GG16 cells as previously described were isolated from GB surgical samples [38]. Genetically modified variants GG16-WT control(ctr) and GG16-PERK-KO were described before [39]. Cells were cultured as GSC enriched neurospheres in NBM with 2% B27 supplement, 20 ng/ml bFGF, 20 ng/ml EGF and 1% L-glutamine (All purchased from Sigma-Aldrich, Darmstadt, Germany). When indicated cells were treated with the chemical PERK inhibitor GSK2606414 (GSK414) (TOCRIS, 5107, UK) at 1µM or Latrunculin B (F-Actin inhibitor) (Sigma Aldrich, Darmstadt, Germany) at 5µM in culture medium. Cells were maintained in an incubator with 5% CO_2_ at 37 °C. For 2D cell culture on hydrogels, first 10 µl of the gels with different stiffnesses were added into the wells of a µ-Slide Angiogenesis chip (Ibidi GmbH, 81506, Germany) and after neurospheres dissociation with accutase (Merk, Germany), 5000 cells in 20 µl NBM^+^ were seeded on top of each hydrogel. After 6 hr in an incubator at 37 °C to allow cell adhesion, 30 µl NBM^+^ was added to each well and cell culturing was prolonged for 7 days for further analyses. Cells were regularly tested for mycoplasma and authenticated by STR profiling.

### Immunofluorescent staining and microscopy

For immunofluorescent (IF) staining, cells on hydrogels were subsequently washed 3 times with PBS for 5 min, fixed with 4% paraformaldehyde (PFA) solution in PBS for 30 min, washed 3 times with PBS and permeabilized with 0.5% Triton-x100 in PBS for 15 min. After 3 times washing with PBS cells were incubated with blocking solution (1% BSA and 3% goat serum in 1x-PBS) for 30 min at room temperature. Following 3 PBS washing steps cells were incubated with the indicated antibodies diluted in a 1% BSA in PBS solution over night at 4 °C. Antibodies/staining used were, beta-Tubulin Monoclonal Antibody (Thermo Fischer Scientific, 22833, Germany) at 1 µg/ml dilution, CoraLite®488-conjugated Vimentin Monoclonal antibody (ProteinTech, 60330, UK) at 1:250 dilution, Alexa-fluor^TM^-Phalloidin (Alexa-594, Thermo Fischer-scientific) at 1:40 dilution, GFAP Monoclonal Antibody (ASTRO6) (Thermo Fischer Scientific, 12023) at 1:200 dilution, recombinant Anti-Vinculin antibody (EPR8185) (abcam, 129002, UK) at 1:600 dilution, Talin Monoclonal Antibody (TA205) (Thermo Fischer Scientific, 28133) at 5 µg/ml dilution, Tensin 1 Polyclonal Antibody (Thermo Fischer Scientific, 116023) at 1:200 dilution, and recombinant Alexa Fluor® 647 Anti-Integrin beta 1 antibody (EPR16895) (abcam, 225270) at 1:100 dilution. Secondary antibody incubation was performed using either Goat anti-Rabbit/mouse IgG Alexa Fluor 488, or 594 secondary antibodies (Thermo Fischer Scientific) at 1:500 dilution in PBS. Finally, after 3 times washing with PBS, cells were mounted with mounting Medium with DAPI^TM^ (ibidi GmbH, 50011, Germany). Microscopic samples were kept at 4 °C until further analyses using the Leica SP8x (Leica Microsystems Co., USA) laser scanning confocal microscope. Images were obtained using the 63x oil immersion lens and intensity of the laser, gain and saturation were kept the same for all the samples to generate comparable data. A scanning depth of 10 µm was used during microscopy to image both cells on top of the gels as well of cells penetrating the gels to ensure accurate imaging.

### Cellular characterization and microscopic data analysis

The Vinculin, Talin, Tensin, Integrin, Tubulin, Vimentin, GFAP and F-Actin surface areas obtained by confocal microscopic imaging were quantified with LAS-X software (Leica Microscopy Co.) and ImageJ software to represent their expression level. Each image was analyzed separately while keeping all process conditions the same. Briefly, original images were switched to 8-bit images. The dimension was corrected in the scale section (set scale). The image was improved by the ImageJ plugins facility to decrease background noise and blurriness of the image, while keeping the color intensity/brightness the same. Using the image threshold, the surface area was marked and measured in µm^2^ and averages from three experiments were used for plotting the data. The surface area was normalized to the cell number.

Cell morphology was evaluated by eye, counting the number of rounded and elongated cells in each microscopic image. In addition, cell shape was also analyzed with LASX software (Leica Microsystem Co.). Briefly, for each individual cell the length (highest measured value) was divided by the width (lowest measured value) in which measured ratios more than 1.5 (≥1.5) representing elongated cells and less than 1.5 (1.5≤X≥1) rounded cells.

### Statistical analyses

Experiments were repeated at least three times unless otherwise indicated. OriginLab (2020b) software was used to plot the data and analyzed with the one-way ANOVA data analyses tool. Data are presented as means with standard deviations (SDs). A significant difference in statistics was considered at *p* < 0.05.

## Results

### PERK mediates stiffness dependent cellular adaptation and differentiation of GSCs

We started by confirming the involvement of PERK in stiffness dependent cellular adaptation of GSCs using the HBP/alginate hydrogels. GG16-WT and GG16-PERK-KO cells were cultured on hydrogels with stiffness of 1, 12 and 35 kPa. Figure 1A shows stiffness dependent F-Actin polymerization in GG16-WT cells, whereas this was not seen in the PERK deficient cells (Figure 1B). F-Actin polymerization of GG16-WT cells increased up to about 5 times from soft to the stiffest matrix, which was not seen in PERK-KO cells that had overall very low levels of F-Actin expression with only minor detectable quantities at the stiffest hydrogels (Figure 1C, E). These patterns of F-Actin polymerization correlated with a change in cell morphology, from more round in soft - to more elongated cells in the stiffer matrices, which depended on the presence of PERK (Figure 1D, F). Since cell elongation often reflects GSC differentiation, we also stained the cells for the astrocytic differentiation marker glial fibrillary acidic protein (GFAP). Indeed, GFAP expression showed a sharp stiffness dependent increase in GG16-WT cells, while GFAP in PERK-KO cells were hardly detected at lower hydrogel stiffness (Figure 1A, B). Quantification of GFAP showed a 6-fold increase of GFAP in GG16-WT cells cultured in soft to stiffest hydrogel, whereas much lower levels were seen in PERK-KO cells, although still an approximate 2.5-fold increase was seen in the stiffest matrix.

**Figure 1.**
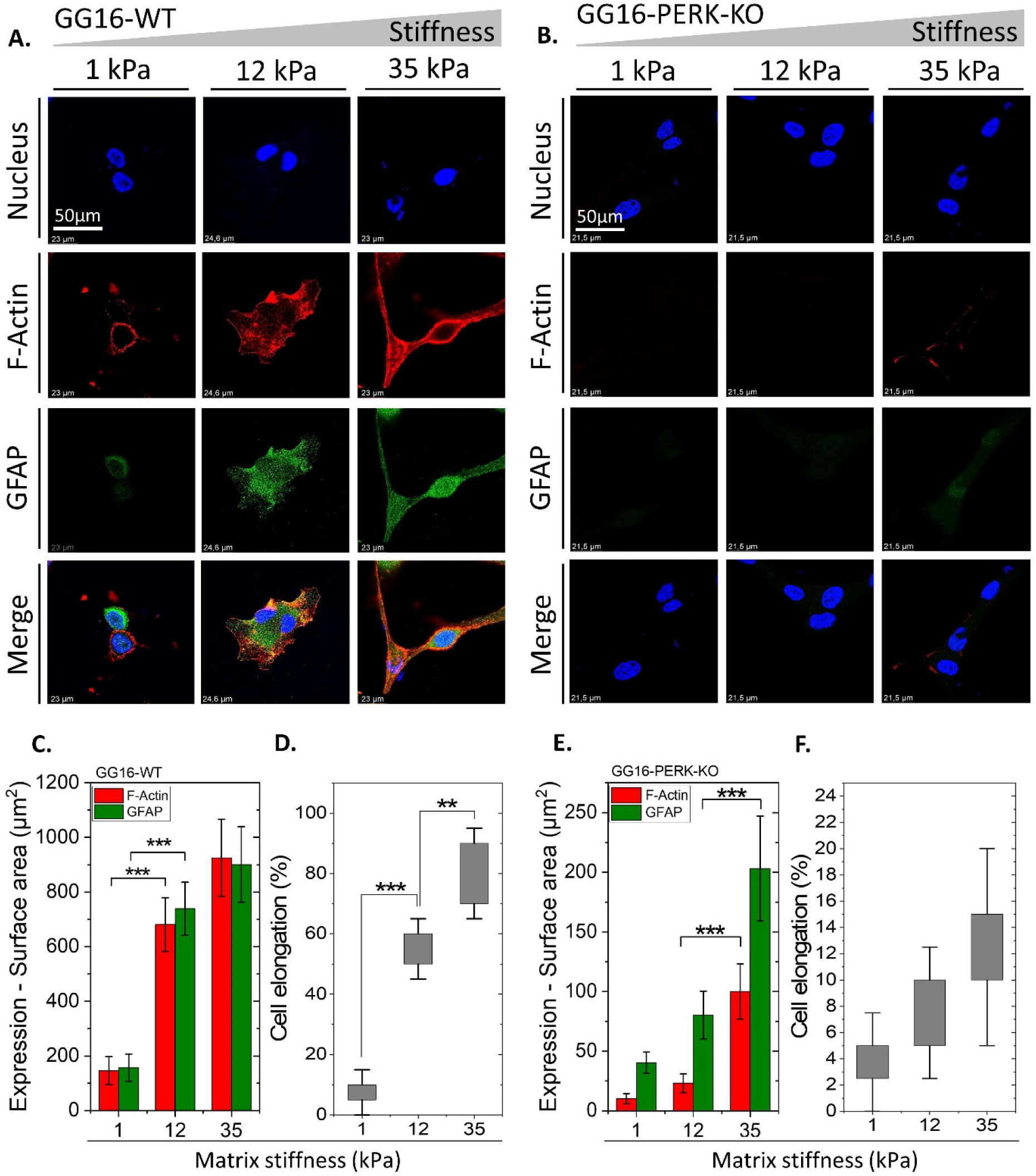
PERK regulates F-Actin polymerization, cytoskeleton remodeling and GSC differentiation. Confocal microscope images of (A) GG16-WT and (B) GG16-PERK-KO cells cultured on increasing matrix stiffness, stained with F-Actin and GFAP (differentiation marker). Expression levels (surface area of staining signals) measured for (C) GG16-WT and (E) GG16-PERK-KO. Cell elongation measured for (D) GG16-WT and (F) GG16-PERK-KO cells. * p ≤ 0.05; ** p ≤ 0.01; *** p ≤ 0.001.

In line with our previous findings, stiffness dependent GG16-WT cell adaptation (morphology, F-Actin polymerization) was hardly affected by exposure to the PERK kinase activity inhibitor GSK414 (Supplementary Figure S1). Stiffness dependent increases in GFAP expression were also independent from PERK kinase activity (Supplementary Figure S1). On the other hand, disruption of F-Actin polymerization by Latrunculin B treatment also impaired stiffness dependent adaptation of GG16-WT cells, and strongly reduced GFAP expression thus mimicking the PERK-KO phenotype (Supplementary Figure S1). As controls, stiffness dependent increases of FLNA and F-Actin seen in WT cells was disrupted in PERK deficient cells (Supplementary Figure S2 and S3), in agreement with our earlier findings [35].

Taken together, stiffness dependent adaptation of GG16 cells also involves the induction of differentiation, which is disrupted in PERK-deficient cells.

### PERK is required for stiffness dependent increases of Vinculin expression

To explore the effect of increasing matrix stiffness on the expression of FAC core proteins known to be involved in mechanotransduction, GG16-WT cells were cultured on 3 increasing hydrogel stiffnesses followed by staining for Talin and Vinculin expression by IF microscopic analyses. Talin binds Integrin complexes to the actin cytoskeleton and Vinculin regulates mechanical signal transmission through this complex upon cell-ECM interactions [32,40]. Vinculin also reinforces the linkage of Talin to Integrins thus initiating a positive feedback loop between the cytoskeleton and FACs [33,41]. Both Talin and Vinculin showed a trend for progressive increased expression upon increasing matrix stiffness (Figure 2A, C). Both proteins showed a cytoplasmic/cell membrane localized punctate pattern, which partially overlapped, and mostly at the cell membrane. Quantification of expression indicated that Vinculin and Talin expression increased from the softest to the stiffest hydrogel by around 1.8 – and 1.4-fold, respectively. Interestingly, in GG16-PERK-KO cells Vinculin expression was much lower, around 6, 8, and almost 11 times lower than in GG16-WT cells, in 1, 12 and 35 kPa hydrogels, respectively (Figure 2B, D). On the other hand, Talin expression was only somewhat reduced in PERK-KO vs WT cells, and the stiffness-dependent increase in expression was similarly to WT cells. These data show that, in the absence of PERK, Vinculin expression is strongly reduced while Talin expression remains comparable to PERK proficient cells upon increasing the matrix stiffness.

**Figure 2.**
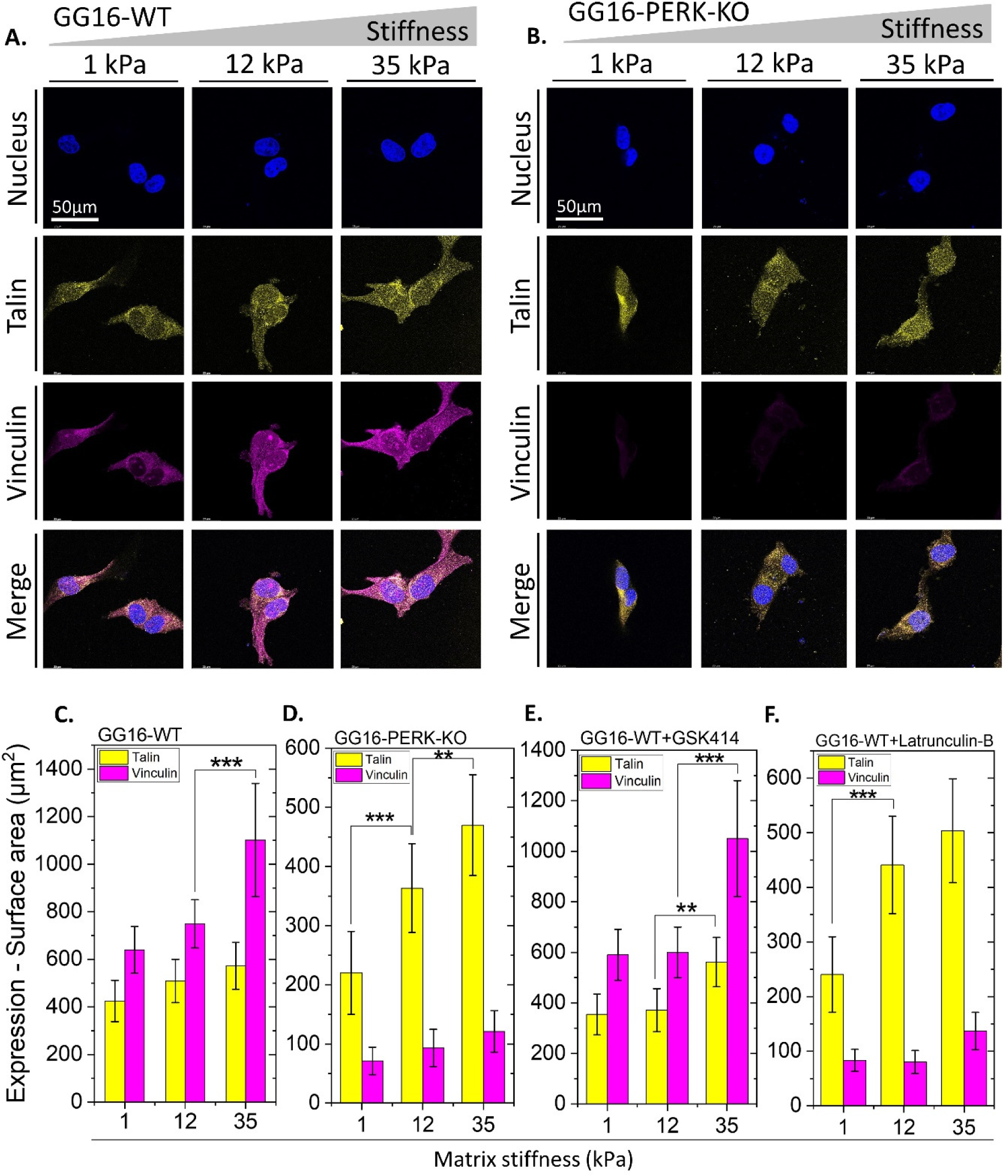
Stiffness dependent upregulation of Vinculin requires presence of PERK and F-Actin remodeling. Confocal microscope images of (A) GG16-WT and (B) GG16-PERK-KO cells cultured on hydrogels with increasing stiffness stained for Talin and Vinculin expression. Quantified Talin and Vinculin expression levels are indicated for (C) GG16-WT, (D) GG16-PERK-KO, (E) GG16-WT treated with GSK414 and (F) GG16-WT treated with Latrunculin B. * p ≤ 0.05; ** p ≤ 0.01; *** p ≤ 0.001.

To examine the involvement of the kinase function of PERK on Talin and Vinculin expression, cells were exposed to PERK kinase inhibitor GSK414. As shown in Figure 2E, GSK414 had no clear effect on both Talin and Vinculin expression showing similar trends as seen in untreated GG16-WT cells (Figure 2C). It indicates that PERK kinase activity is not required for stiffness dependent expression of these FAC components. Next, we examined if F-Actin polymerization was involved in the stiffness dependent increase in Vinculin expression. Therefore, cells were exposed to Latrunculin B, which potently disrupted F-Actin polymerization (see also Supplementary Figure S3 and S4). As quantified in Figure 2F, Latrunculin B treatment sharply decreased Vinculin expression while Talin expression showed a similar pattern as seen in untreated GG16-WT. These results indicate that F-Actin remodeling is also required for stiffness dependent changes in Vinculin expression.

### PERK is required for stiffness dependent increase of Tensin expression

Next, the expression of the FAC compounds Tensin and Integrin-β1 was examined. Integrins are known as the hub of mechanosensing during cell-ECM interactions and actively assist the cells to sense the composition and mechanics of the ECM [21,23,42]. Upon complexation of Integrins and Talin, the recruitment of Tensin reinforces the stability and maturation of FACs. In addition, Tensin stimulates the assembly of additional FACs and promotes their maturation and facilitates cell migration [22,29,40,43]. Both Tensin and Integrin-β1 showed punctuated expression patterns with a trend for stiffness dependent increases in expression (Figure 3A, C). In GG16 cells Tensin expression levels increased around 1.8-fold and Integrin-β1 around 3-fold from soft to the stiffest matrix and colocalization was seen particularly at the cell membrane region (Figure 3C). In contrast, in PERK-KO cells, Tensin expression was hardly detected whereas Integrin-β1 expression was comparable to GG16-WT cells (Figure 3B, D). Together these findings indicate that PERK regulates the increase of Tensin expression upon matrix stiffening while PERK has no effect on Integrin-β1 expression.

**Figure 3.**
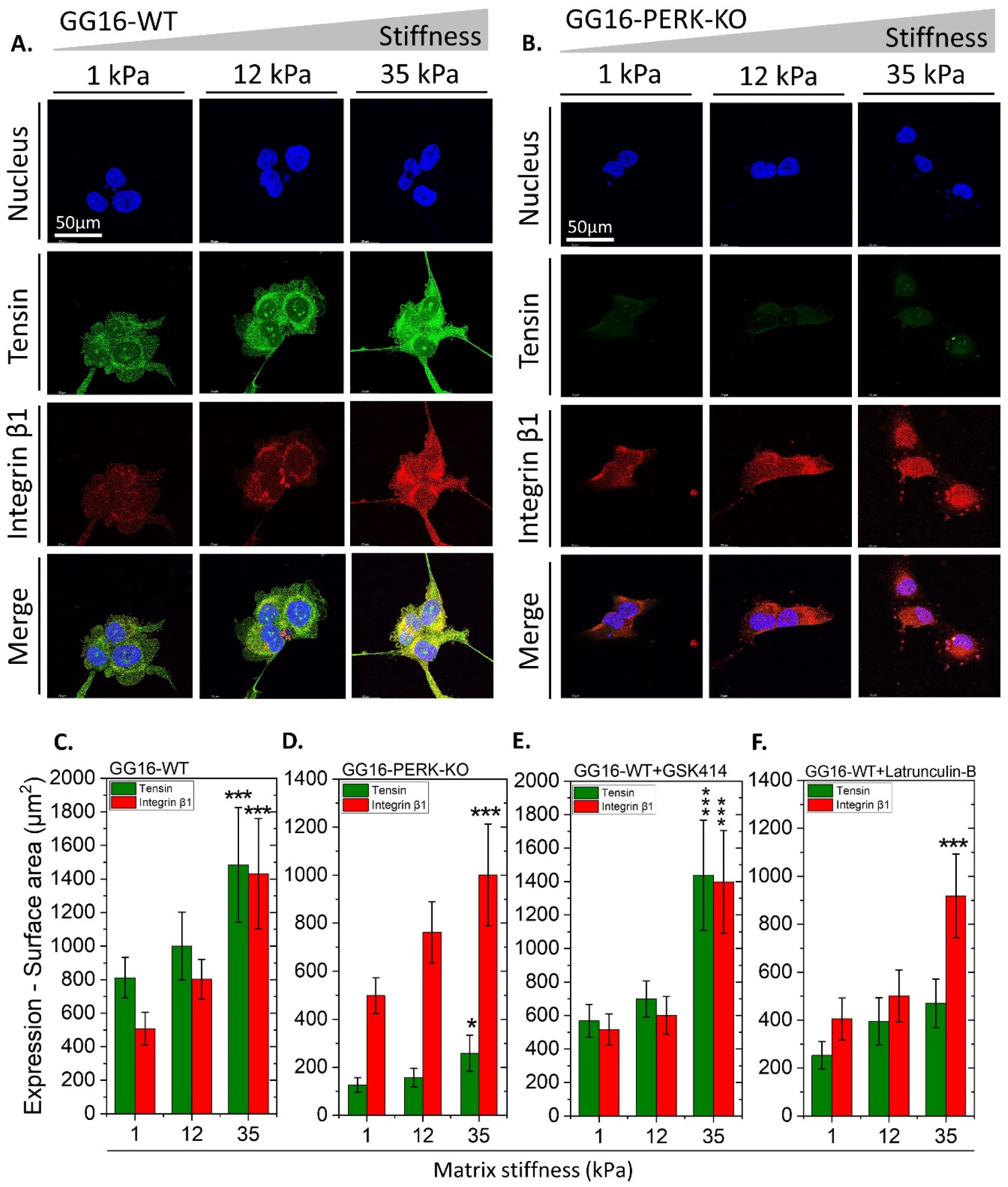
PERK and F-Actin remodeling determine Tensin expression in soft and stiffening matrices. Confocal microscope images of (A) GG16-WT and (B) GG16-PERK-KO cells cultured on increasing stiff matrices and stained for Tensin and Integrin-β1. Tensin and Integrin-β1 expression levels (surface area of staining signals) were measured for (C) GG16-WT, (D) GG16-PERK-KO, (E) GG16-WT treated with GSK414 and (F) GG16-WT treated with Latrunculin B. * p ≤ 0.05; ** p ≤ 0.01; *** p ≤ 0.001.

Next, the involvement of PERK kinase activity on Tensin expression was examined. GSK414 treatment of GG16-WT cells did not have a clear effect on Tensin or Integrin-β1 expression (Figure 3E). At 1 and 12 kPa hydrogels, Tensin and Integrin-β1 showed somewhat lower expression levels in GSK414 exposed cells compared to untreated cells, yet expression increased sharply at 35 kPa under both conditions. Inhibition of F-Actin polymerization by Latrunculin B generated a similar phenotype as seen in the PERK deficient cells (Figure 3F and supplementary Figure S5). These results indicate that F-Actin polymerization has particularly an impact on Tensin expression.

### Loss of PERK results reduces Vimentin expression, increases Tubulin expression and induces an F-Actin/Tubulin switch

FACs are not only connected to the F-Actin network but also components such as Integrins, Talin, and Vinculin can directly interact with Tubulin and Vimentin, facilitating force propagation and mechanical signal transduction via microtubule and intermediate filament networks. To examine the possible involvement of Tubulin and Vimentin in stiffness adaptation, GG16 cells were cultured on the different hydrogels. As shown in Figure 4A and B, GG16-WT and PERK-KO cells showed opposite Tubulin and Vimentin expression patterns. GG16-WT cells showed high expression of Vimentin, which further elevated (around 2-fold) with increasing matrix stiffness, while Tubulin expression was very low (Figure 4C). In contrast, PERK deficient cells had high Tubulin expression levels that peaked at 12 kPa stiffness and dropped at 35 kPa, whereas Vimentin expression was very low and increased only at the stiffest matrix (Figure 4D).

**Figure 4.**
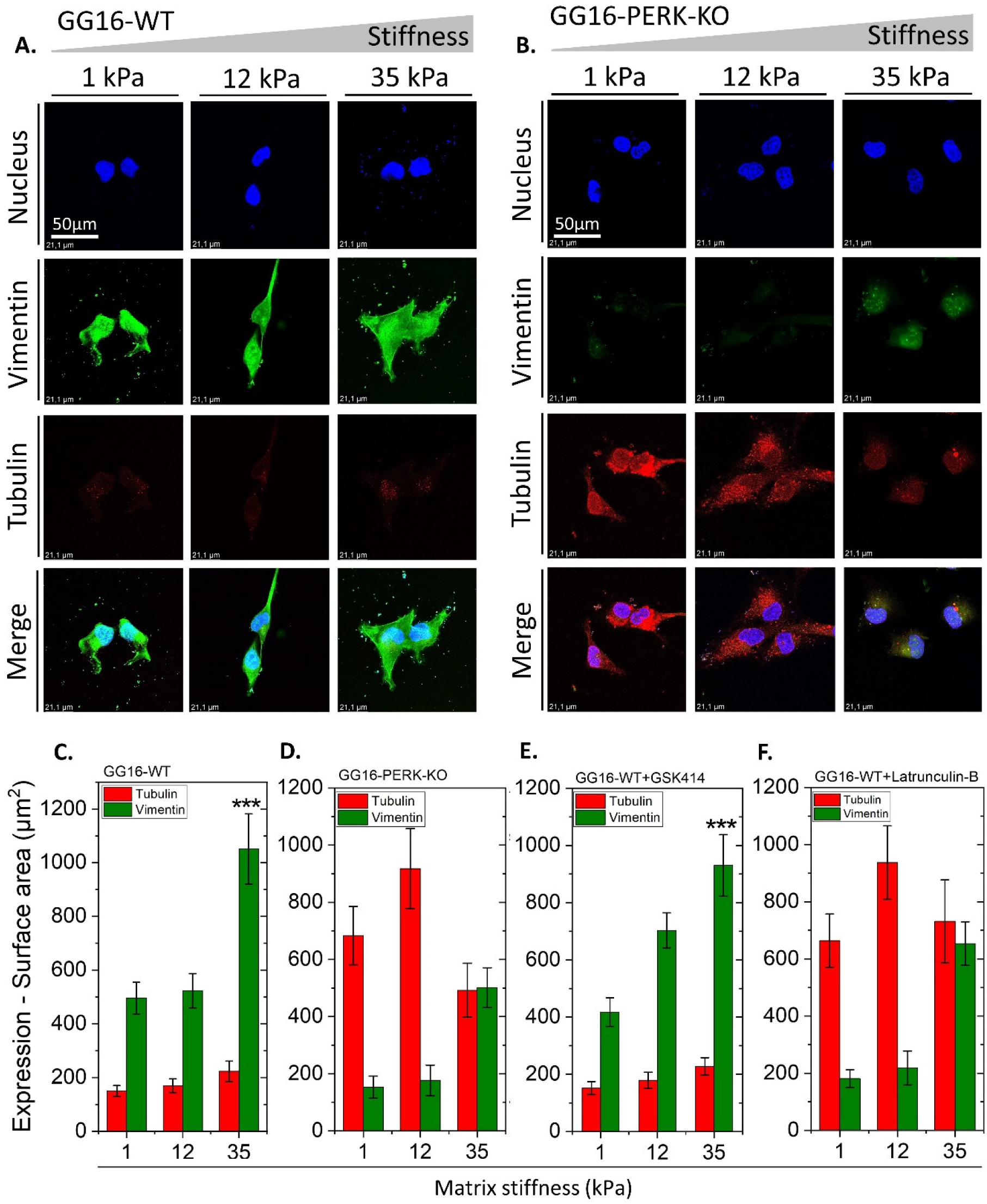
Loss of PERK results in reduced Vimentin and increased Tubulin expression. Confocal microscope images of (A) GG16-WT and (B) GG16-PERK-KO cells cultured on increasing stiff matrices stained for Tubulin and Vimentin. Tubulin and Vimentin expression levels (surface area of staining signals) measured for (C) GG16-WT, (D) GG16-PERK-KO, (E) GG16-WT treated with GSK414 and (F) GG16-WT treated with Latrunculin B. * p ≤ 0.05; ** p ≤ 0.01; *** p ≤ 0.001.

The exposure of GSK414 did not affect Tubulin and Vimentin expression in GG16-WT cells grown on the different matrices when compared to untreated cells (Figure 4C, E). Interestingly, Latrunculin B treated GG16-WT cells showed a similar expression pattern as seen in PERK-KO cells, with reduced Vimentin expression and strongly increased Tubulin expression that also showed a peak at 12 kDa (Figure 4D, F and supplementary Figure S6).

The finding that inhibition of F-Actin polymerization reduced Vimentin expression while Tubulin expression increased sharply suggested that the tubulin network may compensate for loss of the actin network. To examine this further we compared F-Actin and Tubulin expression in GG16-WT and PERK-KO cells. Figure 5A and B shows opposite trends for F-Actin and Tubulin expression in the presence and absence of PERK. GG16-WT cells showed stiffness dependent increases in F-Actin polymerization and a low expression of Tubulin that only slightly increases at the stiffest matrix. In contrast, PERK deficient cells showed strongly impaired F-Actin polymerization, while Tubulin expression is already high in the soft matrix and peaking at the 12 kPa matrix (Figure 5D). Around 12 to 14 times higher Tubulin levels are detected in PERK-KO compared to PERK-WT cells. Interestingly, in PERK deficient cells, Tubulin expression decreased in the stiffest matrix, while F-Actin levels increased (Figure 5D), suggesting that when the F-Actin network is present there is less requirement for the Tubulin network.

**Figure 5.**
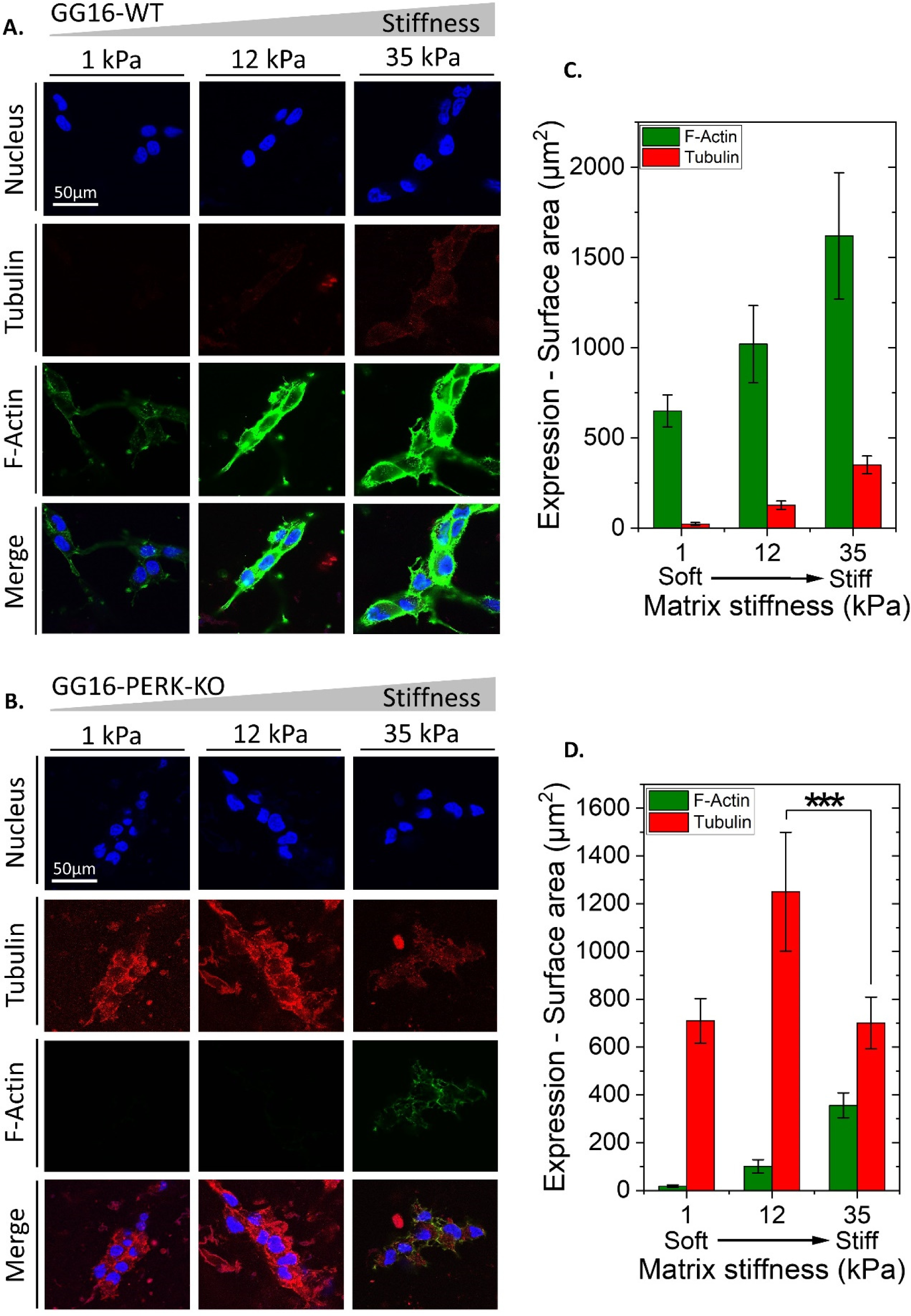
In absence of PERK opposite Tubulin and F-Actin expression levels are seen upon increasing matrix stiffness suggesting a F-Actin/Tubulin compensatory mechanism. Confocal microscope images of (A) GG16-WT and (B) GG16-PERK-KO cells cultured on stiffening matrices stained for F-Actin and Tubulin. Expression levels (surface area of staining signals) measured for (C) GG16-WT and (D) GG16-PERK-KO cells. * p ≤ 0.05; ** p ≤ 0.01; *** p ≤ 0.001.

## Discussion

The current study identified a novel function for PERK in regulating the maturation of FACs in GSCs during the adaptation to an increasing matrix stiffness. Interestingly, the cellular adaptation of GSCs was accompanied by stiffness dependent induction of differentiation that required an increase in F-Actin polymerization. Particularly, the expression of Vinculin and Tensin were regulated by PERK, whereas Talin and Integrin-B1 expression were not affected by PERK. In PERK-deficient cells, reduced Vinculin and Tensin levels could be linked with a disrupted F-Actin polymerization, since similar changes in expression were seen in WT cells treated with an F-Actin inhibitor. Furthermore, Vimentin and Tubulin expression were increased at lower matrix stiffnesses in PERK deficient cells compared to WT cells, suggestive of compensatory activity of the tubulin network. Overall, our findings indicate that the previously identified PERK/FLNA pathway is also involved in regulating force transmission from the ECM to the actin cytoskeleton via FACs at matrix stiffnesses that are representative for normal brain and glioblastoma.

Our finding that the adaptation of GSCs to increasing matrix stiffness is also accompanied by increased expression of the differentiation marker GFAP, which is not observed in PERK deficient cells, links PERK function to GSC differentiation. This finding is in line with our previous work, showing that PERK deficient GSCs compared to WT cells display aberrant differentiation that is characterized by diminished cell adhesion, higher levels of SOX2 and lower levels of GFAP [39]. It should be noted that the previous studies were performed on regular cell culture plastics (high stiffness), indicating that PERK regulates differentiation of GSCs at a broad stiffness range. The found stiffness dependent induction of differentiation of GSCs is in accordance with previous studies reporting that stem cell differentiation is regulated by ECM stiffness, such as in mesenchymal - and muscle stem cells [44,45].

The activity of FACs are strongly influenced by matrix stiffening and orchestrate cytoskeleton remodeling and subsequent adaptive responses [46,47]. FACs are also involved in tuning the mode of actin flow, where they mediate cell migration via actin retro-grade flow [23,48]. In our study, integrin-β1 and Talin expression, representing an early stage of FAC formation, could be detected at low stiffness conditions. Integrins are the main contact point to the ECM for the cells to sense mechanical cues, as was shown before [23,34,49]. Talin links the integrins to the cytoskeleton and forms the backbone of FAC formation [22]. PERK did not influence the formation of the Integrin-β1/Talin complex in GSCs, which is considered as a hub of mechanosensing. However, PERK strongly regulated the expression of both Vinculin and Tensin upon matrix stiffening. Vinculin is a main player in the maturation of the FAC and acts as an signal carrier between Integrin-β1/Talin and the F-Actin cytoskeleton [22,32,50,51]. Tensin also plays a significant role in the maturation of FACs by reinforcing Vinculin binding and provides stability to the F-Actin filaments interactions within the FACs [21,22]. In agreement with their F-Actin binding activity, impairing F-Actin polymerization in the GSCs also reduced Vinculin and Tensin levels. Of note, FLNA levels also increased in a stiffness dependent way and is known to bind and stabilize the F-Actin cytoskeleton and in the anchoring of membrane proteins [52]. PERK may thus regulate the formation of FACs through FLNA and or indirectly by F-Actin remodeling.

The intermediate filament protein, Vimentin, was expressed at high level in GSCs at all matrix stiffnesses tested. Vimentin is known to bind Integrins, Talin, and Vinculin, and also interacts with F-Actin and FLNA, thus reinforcing the F-Actin network to facilitate cytoskeleton remodeling [53–57]. However, Vimentin expression was sharply decreased in PERK deficient GSCs, which could be associated to the inhibition of F-Action polymerization. Previously, FLNA was shown to regulate Vimentin expression and functioning [56], suggesting that PERK regulates Vimentin via FLNA or indirectly via impaired F-Actin polymerization. The precise mechanism by which PERK regulates these components of FAC maturation remains to be elucidated.

Furthermore, we unveiled a potential rescue mechanism for F-Actin network impairment in PERK-deficient cells. In the absence of PERK, Tubulin expression was increased and Tubulin also has binding sites to Integrin complexes and Talin. Although Tubulin also can interconnect to some extent with F-Actin and FLNA, it is largely functioning independently [46,58–60]. In PERK deficient cells, the Tubulin network may compensate for impaired F-Actin remodeling and connect to Integrin/Talin complexes. Figure 6 summarizes the main findings of the current study.

**Figure 6.**
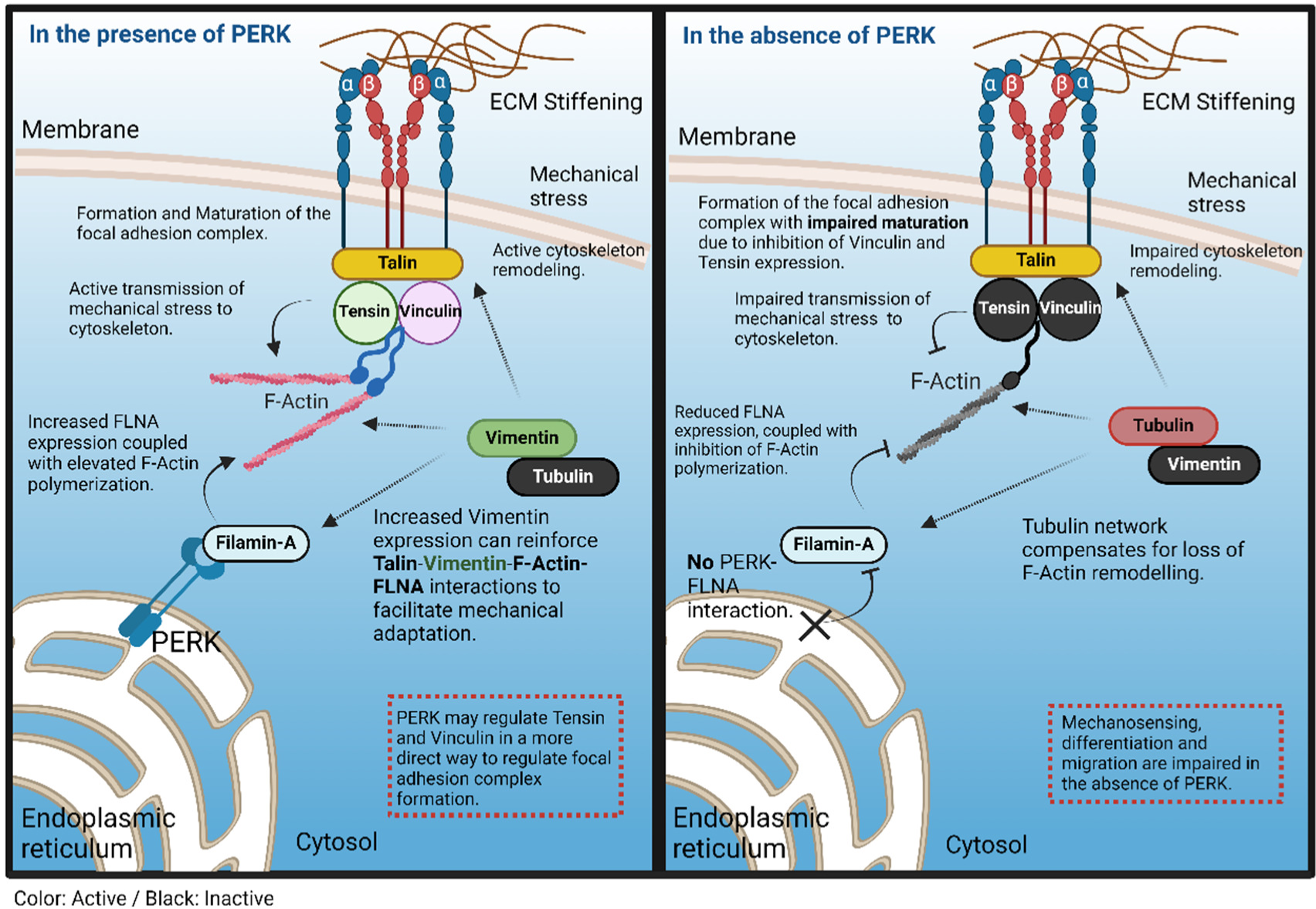
Model of PERK dependent regulation of focal adhesion complex. The maturation of the focal adhesion complex (FAC) and force transmission from the ECM to the cytoskeleton is schematically depicted in the presence or absence of PERK. PERK could affect FAC maturation indirectly via regulating F-Actin remodeling via PERK/FLNA interactions and subsequent recruitment of Tensin and Vinculin, or through an as yet unidentified mechanism by which PERK regulates Tensin and Vinculin expression and association with the FAC. GSCs still sense and adapt to mechanical stress in the absence of PERK through the tubulin cytoskeleton that may provide a rescue mechanism for impaired F-Actin remodeling.

## Conclusion

In conclusion, we found a new non-canonical function of PERK in the regulation of FAC formation during the physiological stiffening of the ECM in glioblastoma that was mimicked by stiffness tunable hydrogels. PERK could indirectly regulate FAC formation via FLNA/F-Actin or in a more direct way by regulating the expression of Tensin and Vinculin. The precise molecular mechanism remains to be elucidated as well as the possible targetability of PERK, independent of its kinase function, for therapeutic purposes.

## Author contributions

Frank Kruyt conceptualized the study. Frank Kruyt, Patrick van Rijn and Mohammad Khoonkari designed the study. Mohammad Khoonkari and Dong Liang performed the experiments, collected and analyzed data. Frank Kruyt, Patrick van Rijn and Marleen Kamperman supervised the project. Mohammad Khoonkari, Frank Kruyt drafted the original paper. Patrick van Rijn and Marleen Kamperman reviewed the paper. Frank Kruyt, Patrick van Rijn and Marleen Kamperman arranged the funding.

## Acknowledgments

Mohammad Khoonkari was financially supported by the Zernike Institute for Advanced Materials at the University of Groningen, including funding from the Bonus Incentive Scheme (of the Dutch Ministry for Education, Culture and Science (OCW)). Dong Liang was supported by the China Scholarship Council [201908320416] and the University of Groningen.

## Conflict of Interest Statement

P.v.R. also is co-founder, scientific advisor, and share-holder of BiomACS BV, a biomedical-oriented screening company. The other authors declare no other competing interests.

## Data Availability Statement

Not applicable.

