## Supplementary figures for "The Unfolded Protein Response Sensor PERK Mediates Mechanical Stress-induced Maturation of Focal Adhesion Complexes in Glioblastoma Cells"

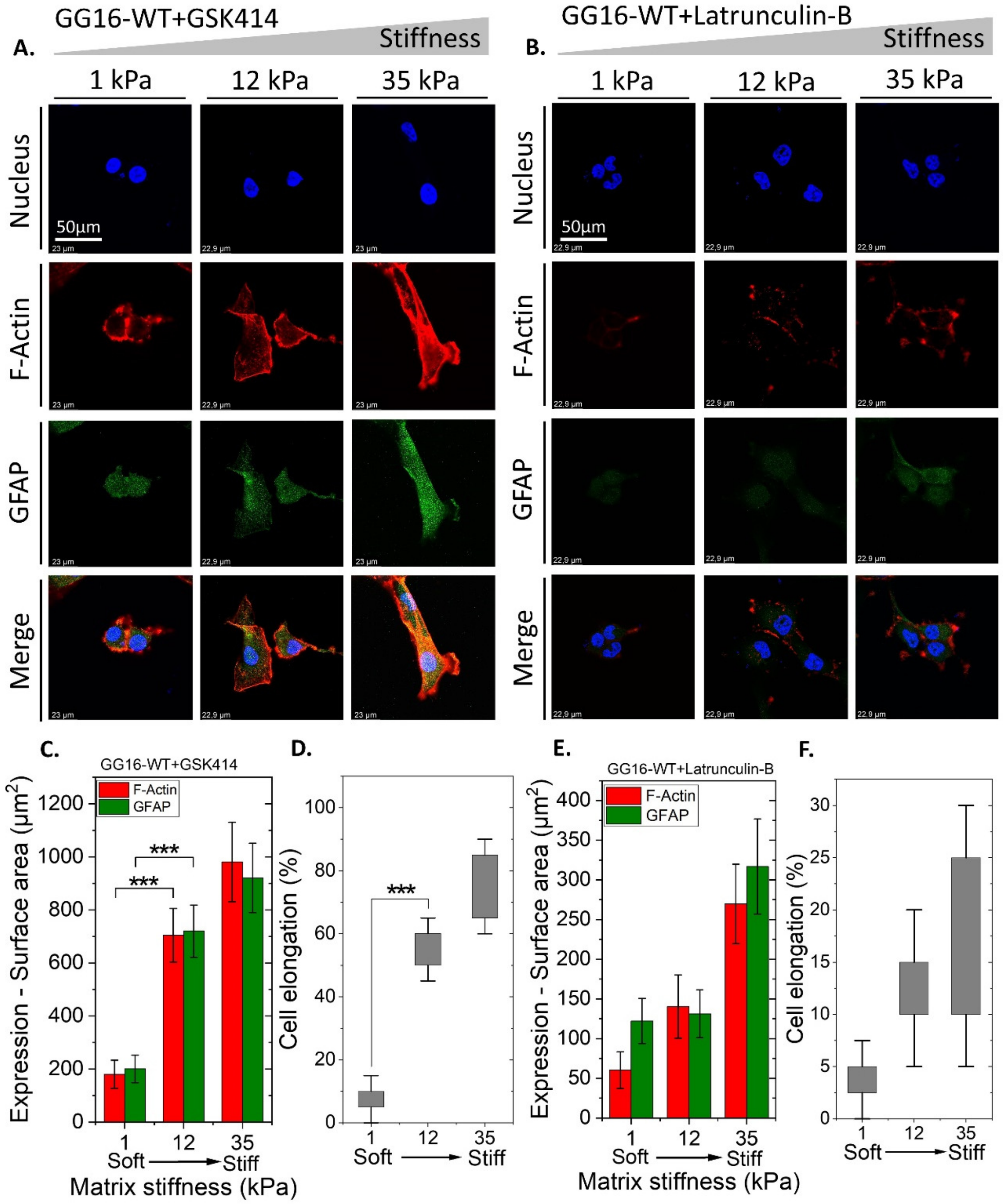


**Figure S1.** Confocal microscope images of GG16-WT cells treated with (A) GSK414 and (B) Latrunculin B and cultured on different matrix stiffnesses stained for F-Actin and GFAP . Expression levels (surface area of staining signals) measured for (C) GG16-WT+GSK414 and (E) GG16-WT+Latrunculin B. Cell elongation measured for (D) GG16-WT+GSK414 and (F) GG16-WT+Latrunculin B.

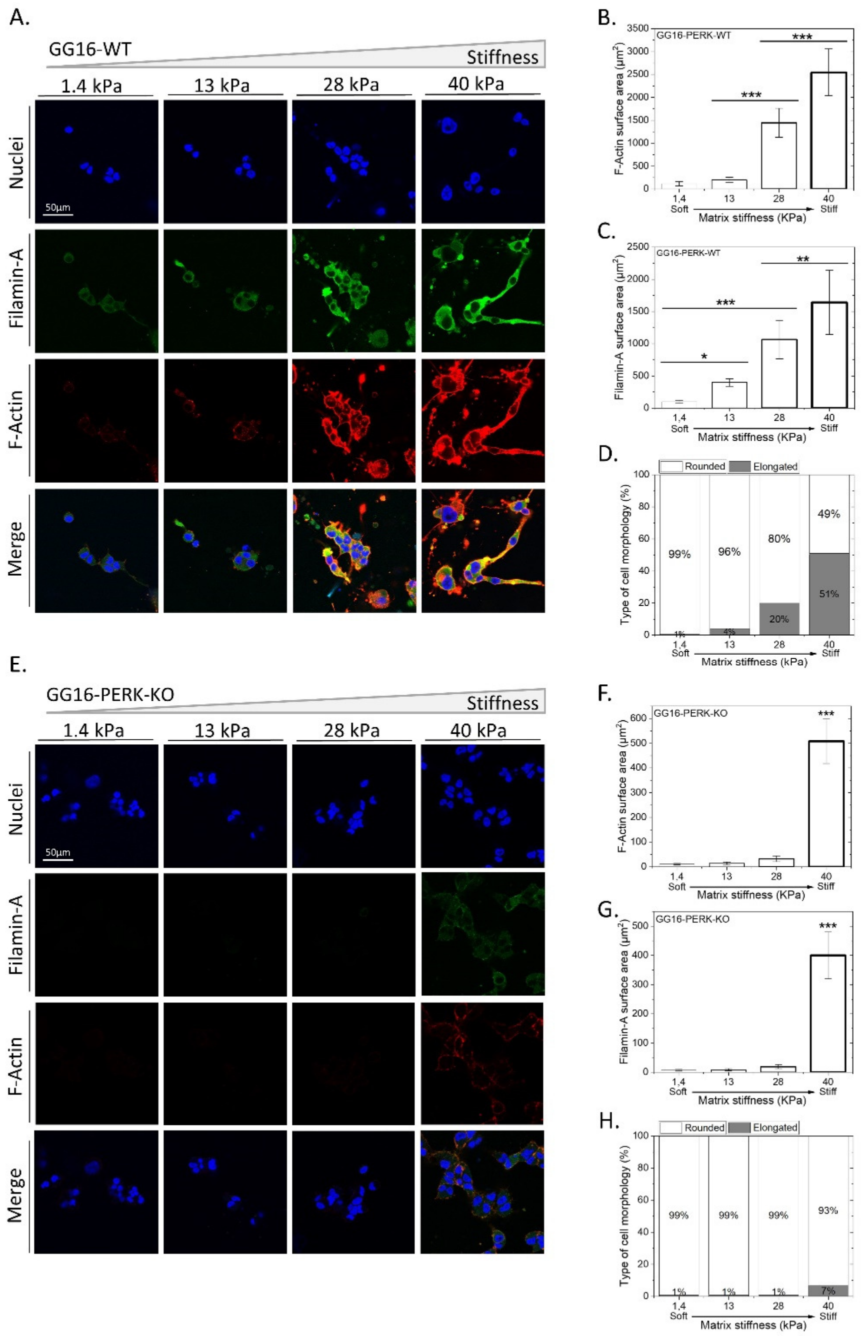


**Figure S2.** PERK deficient GG16 cells are impaired in cellular adaptation to increasing stiffness that is linked with aberrant FLNA expression. (A, E) GG16-WT and PERK-KO cells were cultured for 6 days on hydrogels with different stiffness and stained with DapiTM (Blue), Alexa fluorTM 546-Phalloidin (Red) and FLNA - Alexa fluorTM 488 (Green). Cell morphology, F-Actin and FLNA expression was quantified and is depicted in (B-D) and (F-H) for GG16-WT and PERK-KO cells, respectively. Cell morphology (from round to elongated), F-Actin and FLNA expression altered gradually in a stiffness-dependent manner in GG16-WT cells which was not seen in PERK deficient cells.

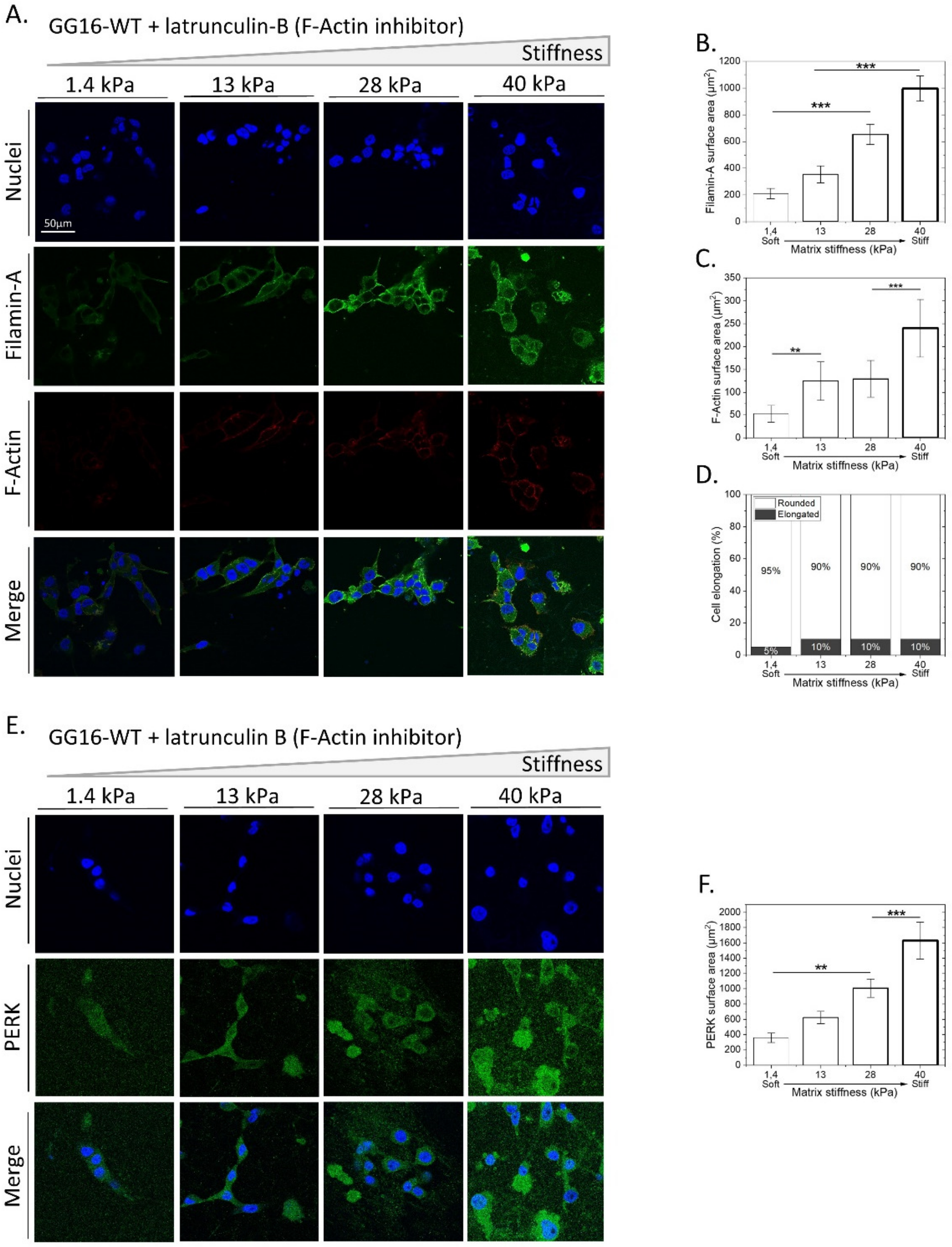


**Figure S3.** Inhibition of F-Actin polymerization mimics phenotype of PERK deficient cells by impairing cellular adaption to matrix stiffness. (A, E) GG16-WT cells treated with Latrunculin B were cultured for 6 days on hydrogels of varying stiffness. Cells were stained for F-Actin, FLNA and PERK  and nuclei and imaged with confocal microscopy. Specific fluorescence was quantified for FLNA (B), F-Actin (C), cell elongation (D) and PERK expression (F).

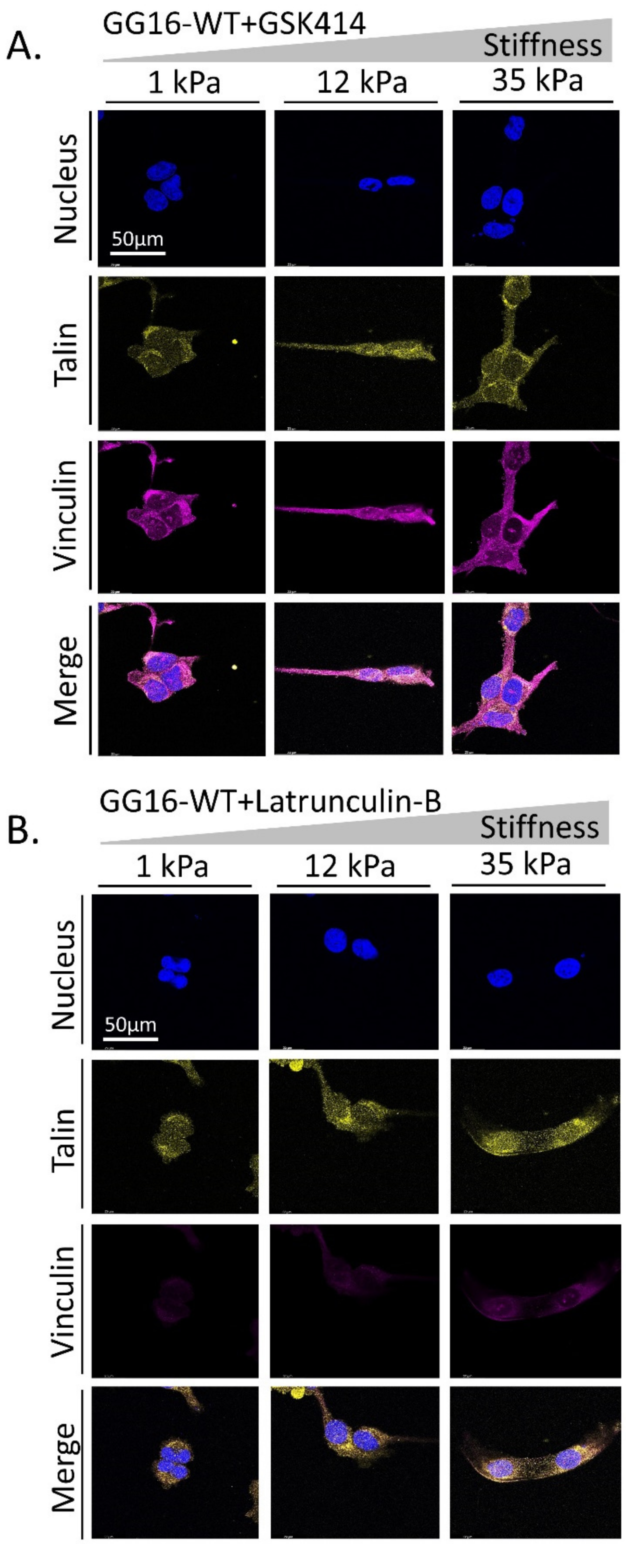


**Figure S4.** Confocal microscope images of GG16-WT cells treated with (A) GSK414 and (B) Latrunculin B, cultured on a hydrogels with indicated matrix stiffness and stained for Talin and Vinculin.

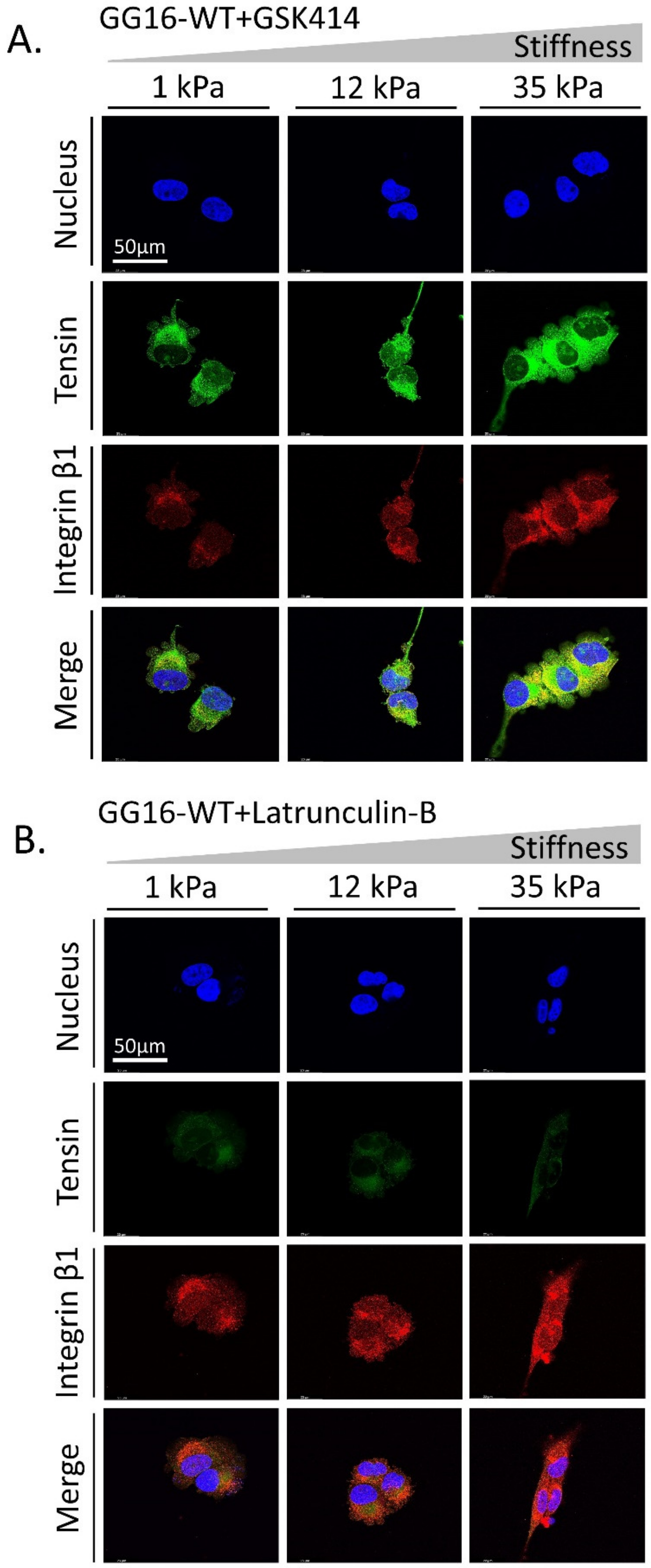


**Figure S5.** Confocal microscope images of GG16-WT cells treated with (A) GSK414 and (B) Latrunculin B and cultured at different matrix stiffnesses and stained for Tensin and Integrinβ1 expression.

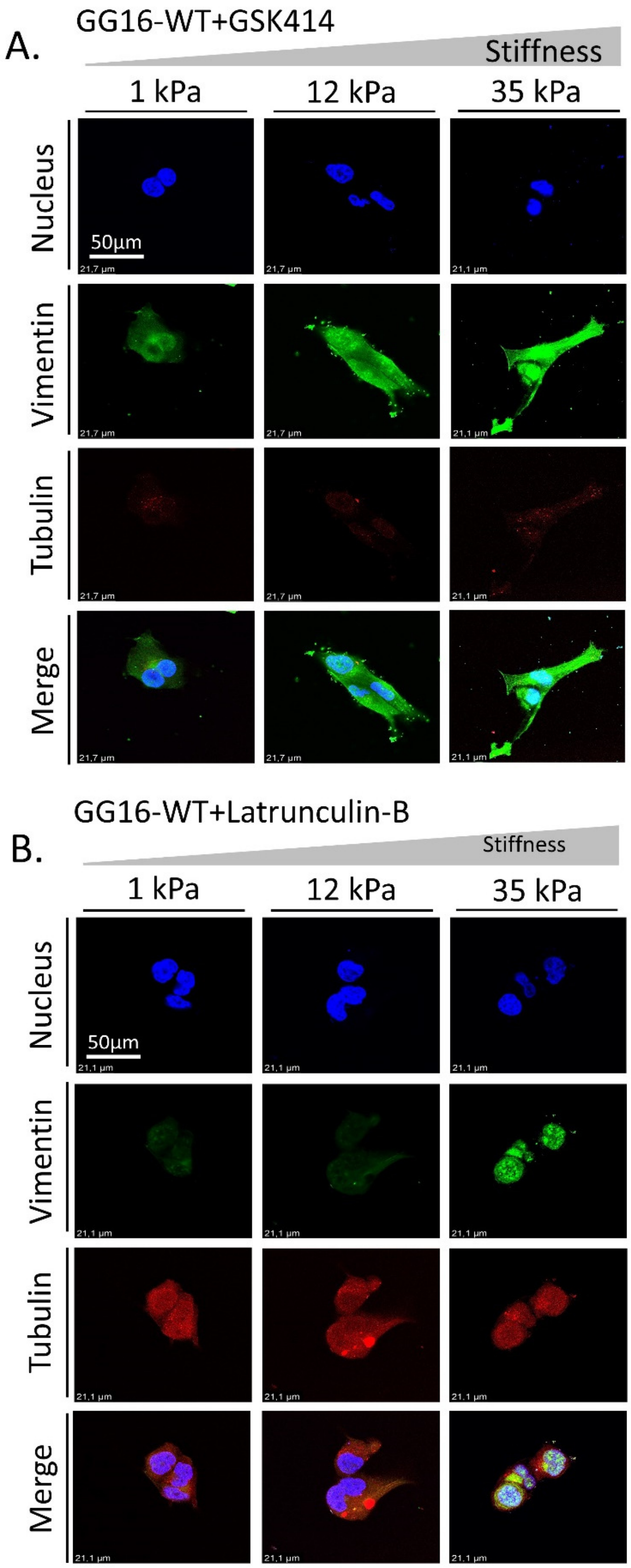


**Figure S6.** Confocal microscope images of GG16-WT cells treated with (A) GSK414 and (B) Latrunculin B and cultured on a range of matrix stiffnesses stained for Tubulin and Vimentin expression.
